# Coding of interceptive saccades in parietal cortex of macaque monkeys

**DOI:** 10.1101/2021.07.05.451092

**Authors:** Jan Churan, Andre Kaminiarz, Jakob C. B. Schwenk, Frank Bremmer

**Author notes:** Corresponding author: Jan Churan.

## Abstract

The oculomotor system can initiate remarkably accurate saccades towards moving targets (interceptive saccades) the processing of which is still under debate. The generation of these saccades requires the oculomotor centers to have information about the motion parameters of the target that then must be extrapolated to bridge the inherent processing delays. We investigated to what degree the information about motion of a saccade target is available in the lateral intra-parietal area (area LIP) of macaque monkeys for generation of accurate interceptive saccades. When a multi-layer neural network was trained based on neural discharges from area LIP around the time of saccades towards stationary targets it was also able to predict the end points of saccades directed towards moving targets. This prediction, however, lagged behind the actual post-saccadic position of the moving target by ^~^80 ms when the whole neuronal sample of 105 neurons was used. We further found that single neurons differentially code for the motion of the target. Selecting neurons with the strongest representation of target motion reduced this lag to ^~^30 ms which represents the position of the moving target approximately at the onset of the interceptive saccade. We conclude that - similarly to recent findings from the Superior Colliculus (Goffart et al., 2017) – there is a continuum of contributions of individual LIP neurons to the accuracy of interceptive saccades. A contribution of other gaze control centers (like the cerebellum or the frontal eye field) that further increase the saccadic accuracy is, however, likely.

## Introduction

Visual tracking of a moving object is achieved by combinations of saccades that bring the image of the object of interest onto the fovea and smooth pursuit eye movements that slowly follow a once foveated target. Saccades towards moving targets were shown to be remarkably accurate (Fuchs, 1967; Casanello et al., 2008; Fleuriet and Goffart, 2012). This performance of the saccadic system is striking since it must be able to extrapolate the motion trajectory of the target to account for its own processing time before saccade onset (typically 100 to 300 ms) as well as for the duration of the saccade (typically 20 to 50 ms) to accurately match the post-saccadic eye position with the position of the moving object. Which brain areas contribute to the processing and extrapolation of the target motion for purposes of interceptive saccades is still under discussion. It was previously shown that the activity of neurons in the superior colliculus (SC) does not account for the component of a saccade vector that is caused by the motion of a saccade target (Keller et al., 1996). The SC neurons in that study shifted their motion fields dependent on whether the saccades were made to stationary or moving targets in a manner that suggested that they only reflect the location of the appearance of a saccade target but not its subsequent motion. This finding gave rise to the so called ‘dual-drive’ hypothesis (Optican and Quaia, 2002; Guan et al., 2005; Optican and Pretegiani, 2017) stating that the motion information is processed in parallel and is added to the motor command at a late stage of processing probably by the caudal fastigial nuclei of the cerebellum. This notion was later refined as Goffart et al. (2017) have shown that SC-neurons to a differential degree participate in generation of the motion related component of interceptive saccades. Beside these findings about the role of sub-cortical areas in processing of interceptive saccades, cortical contributions along the dorsal visual pathway are little investigated yet. Lesions in the middle temporal area (MT) of macaque monkeys, which is a major center in perception of visual motion, were shown to selectively reduce the accuracy of interceptive saccades while leaving saccades to stationary targets unimpaired (Newsome et al. 1985). Target motion related signals were also reported in the Frontal eye-field (Barborica and Ferrera, 2003; Xiao et al., 2007; Ferrera and Barborica, 2010) that receives input from the cortical motion areas (Tian and Lynch, 1996) and has direct efferent connections to the SC (Segraves and Goldberg, 1987; Stanton et al., 1988) and to the oculomotor areas in the brain stem (Schiller et al., 1980).

The lateral intraparietal area (LIP) is a major processing stage in the dorsal visual pathway as well as in the saccade generating circuit. Its connections make it well suited for processing of moving targets. It receives strong inputs from areas MT and the medial superior temporal area (MST, Blatt et al., 1990) that are almost exclusively engaged with processing of visual motion and sends efferent projections to the SC (Paré and Wurtz, 2001) and the frontal eye field (FEF, Schall et al., 1995) which are the major sources of saccadic motor commands. Results from an earlier investigation (Bremmer at al., 2016) have shown that LIP neurons carry information about the motion of a saccade target. The goal of the current study was to specify to what degree information provided by LIP neurons can contribute to the accurate execution of interceptive saccades in 2D oculomotor space. To this end, we trained a multi-layer model to reproduce the trajectories of saccades to stationary targets from the perisaccadic activity of a sample of LIP neurons. Once this model was established, we used activities from the same neurons collected during interceptive saccades as inputs to the model and tested to what degree they account for the interceptive component of the saccade trajectory, that is, the deflection of the saccade end point in the direction of target motion.

## Methods

### Physiological preparation

Two adult male monkeys (macaca mulatta) participated in the study. In two separate surgeries each monkey was implanted with a head holder and a recording chamber. Based on MRI scans prior to surgery, the chamber (inner diameter 14 mm) was centered at a position 3 mm posterior from the interaural line and 15 mm lateral from the longitudinal fissure (P3/L15) to access the region of the intraparietal sulcus (right hemisphere in both monkeys). During the experiments, the correct position of the electrode within the lateral intraparietal area (LIP) was determined based on properties of the neurons under study as well as properties of neurons in the neighboring cortical areas, predominantly the ventral intraparietal area VIP in the depth of the intraparietal sulcus, which was identified by directional selectivity for visual motion. The correct position of the recordings was later confirmed by histology in one of the monkeys (O). The other monkey (S) still participates in ongoing recordings. All procedures had been approved by the regional authorities and were in accordance with the published guidelines on the use of animals in research (European Communities Council Directive 2010/63/EU).

Single-unit recordings were done using standard tungsten microelectrodes (FHC, Bowdoin, USA) with an impedance of ^~^2 MΩ at 1 kHz that were positioned by a hydraulic micromanipulator (MO-95, Narishige, Tokyo, Japan). A stainless-steel guiding tube was used for transdural penetration and support of the electrode. The neuronal signal was processed using a commercial system (Alpha Omega, Nof HaGalil, Israel). It was band-pass filtered (cut-off frequencies at 500 Hz and 8000 Hz) and sampled at 44 kHz.

### Apparatus

During recordings, the monkeys were sitting head-fixed in a primate chair in a dark room, and their eye-position was monitored at 1000 Hz using a video-based eye tracker (EyeLink 1000, SR Research, Ottawa, Canada). The chair was positioned at a distance of 97 cm from a semi-transparent screen (size 160 cm x 90 cm, subtending the central 79 deg x 50 deg of the visual field) on which the visual stimuli were back-projected using a PROPixx-projector (VPixx Technologies, St-Bruno de Montarville, Canada) running at a resolution of 1920 x 1080 pixels and at a frame rate of 100 Hz.

### Paradigms

The experimental paradigms were implemented using Matlab (R2012a, MathWorks, Natick, USA) and the psychophysics toolbox (Brainard, 1997; Pelli, 1997; Kleiner et al. 2007) on a standard Windows (v7, Microsoft, Redmond, USA) PC (Dell Precision T5810, Round Rock, USA). In all following experiments the saccade targets were always presented on a grey background (luminance 40 cd/m2). Before the main experiment started, the saccadic tuning of the neuron under investigation was tested during visually guided saccades to stationary targets. In this pre-test, saccades with different directions (covering the full fronto-parallel angular space of 360° in steps of 45°) and different amplitudes (typically between 7° and 20°) were used to find a saccade vector yielding strong peri-saccadic responses. This saccade-vector was then used during the main experiment, as the ‘preferred vector’ of the neuron. A saccade vector of the same amplitude but opposite direction (‘anti-preferred vector’) was also used in the main paradigm, although, the (usually very low) peri-saccadic activities from this saccade vector were only utilized to calculate the selectivity of the neuronal responses that was a criterion for a neuron’s inclusion into the main analysis.

In the main paradigm two types of trials were employed, the ‘stationary’ trials in which the monkeys made regular visually guided saccades towards stationary targets at different spatial positions and ‘interceptive’ trials where they made saccades towards targets moving in one of eight directions (Figure 1). At the beginning of each trial the monkey fixated a target (small red square, side length 0.8°) placed at an eccentric position on the screen for a random duration (500 to 700 ms). This initial position of the target was chosen so that saccades towards the locations around the screen center yielded strong peri-saccadic responses (relative to the baseline activity) at least for a part of the investigated target locations. Then, in the ‘stationary’ trials (Figure 1, upper panel) the fixation spot stepped to one of 17 predefined positions placed around the center of the screen and the monkey’s task was to make a visually guided saccade to that position. The positions of the target were chosen in the center of the screen (0°/0°) and along the cardinal as well as the oblique axes in distances of 2° and 4° from the screen center. These specific positions were chosen to be identical with the positions of the moving targets in the interceptive trials after 200 ms and after 400 ms of motion and hence to be close to the average landing points of the interceptive saccades. The saccade had to be initiated within 300 ms from the stimulus step and land within an 8° x 8° window around the target. After the saccade, the monkey had to fixate the target for another 500 ms to obtain a liquid reward. Typically 10 trials were collected for each of the 17 different target positions. ‘Interceptive’ trials (Figure 1, lower panel) started with fixation of a stationary target at the same eccentric position as in the ‘stationary trails’. The monkey then made a saccade towards a target that first jumped to the center of the screen and then immediately started moving into one of 8 directions at a speed of 10°/s. Again, the saccade had to occur within 300 ms after the step of the target and had to land within an 8°x8° window around the current target position. After the saccade the monkey had to follow the target using smooth pursuit eye movements (SPEM) for another 800 ms to receive a liquid reward. Typically, 15 trials were collected for each direction of stimulus motion. Both, the stationary and the interceptive trials were presented randomly interleaved during a single measurement. In the following we will name the saccades to the stationary targets ‘regular saccades’ and those to moving targets ‘interceptive saccades’.

**Figure 1:**
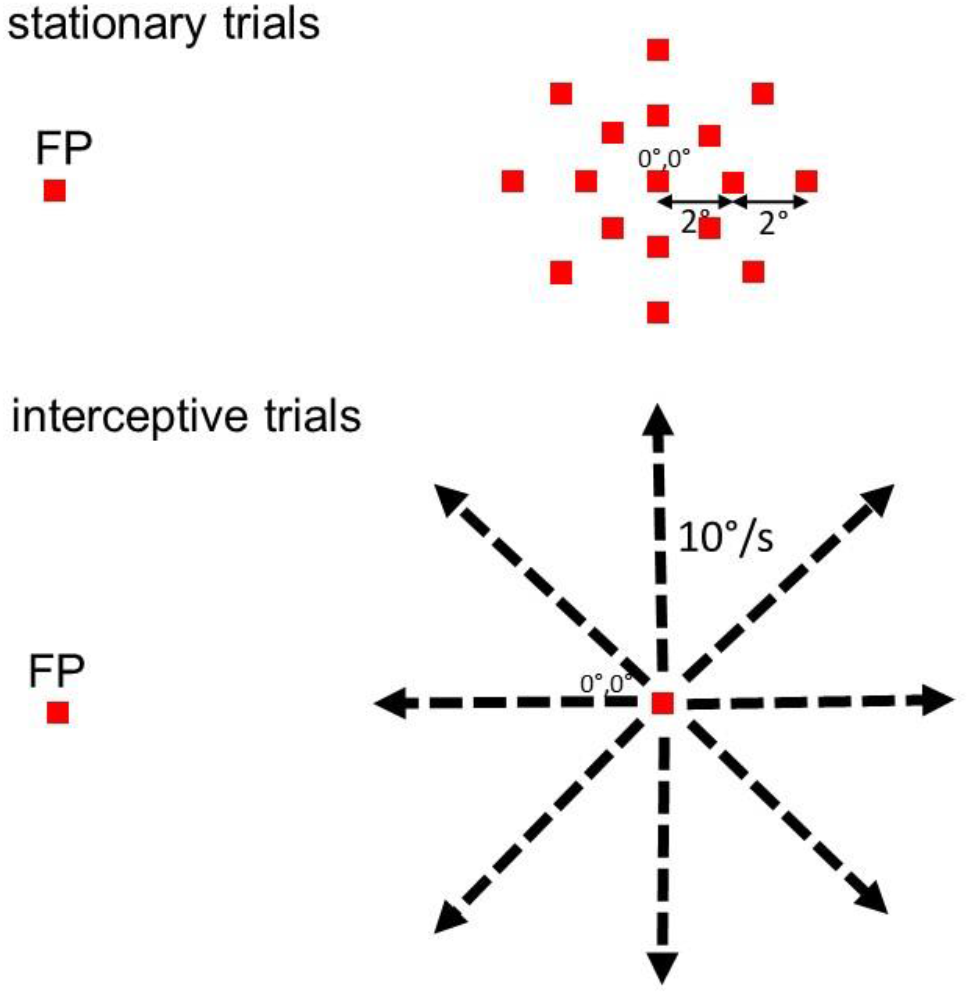
Sketches of the two types of trials that were used in the main paradigm. In the ‘stationary’ trials (upper panel) the monkey had to make a visually guided saccade from an eccentric fixation point (FP) to one of 17 target positions that were presented around the center of the screen. The possible positions of the target are depicted on the right side of the upper panel. They included the center of the screen and positions that were 2° and 4° away from the center in the four cardinal and four oblique directions. These chosen positions represent locations at which the moving target in the ‘interceptive’ trials arrives after 200 ms and 400 ms of motion. In the ‘interceptive’ trials (lower panel) the monkey made a saccade from an eccentric position (which was the same as in the stationary trials) to a target that first stepped to the center of the screen and then immediately moved in one of eight directions at a constant speed of 10°/s.

### Data processing and analysis

#### Processing of eye movements

The eye position signal was smoothed by a moving average filter with a span of 15 ms. Then the initial saccades were detected using a double velocity criterion in which at first time-windows were detected at which the eye-velocity exceeded 100°/s and then in a second step the beginning and the end of a saccade were identified when the velocity before and after these time windows first fell below 30°/s. The pre-saccadic and the post-saccadic eye positions were calculated as averages in a time window 6 - 16 ms before the start and 2 - 4 ms after the end of the saccade. The very short post-saccadic time window was chosen to avoid a contamination by the subsequent SPEM in the interceptive trials. The results were confirmed visually for a subset of the trials to give accurate estimates of pre- and post-saccadic eye positions.

#### Processing of neuronal data

Single units were isolated using a semi-manual spike sorter (Plexon Inc, Dallas, Texas). To this end we used a threshold on the electrode signal that was set manually to separate the action potentials from noise. The samples that exceeded the threshold were further analyzed using principal components as well as other features that were derived from the signal (like local maxima and minima). Then clusters of samples with similar properties were identified visually and grouped together as a single unit. For a detailed description of the sorting process see the offline User Guide (Plexon, 2020). The procedures used for further analysis of the data were written in MATLAB (R2012a, MathWorks, Natick, USA),

##### Generation of a pseudo-population

The data basis for this investigation consists of the activity of a sample of 105 neurons (50 for monkey S, 55 for monkey O) that were tested using both paradigms as described above. All neurons have shown a significant difference in activity (p<0.01, t-test) between the preferred and the anti-preferred saccade direction in a time window between 100 ms before and 100 ms after the saccade onset.

Separate datasets were required for training and validation of the neural network model. Thus the data from the stationary trials was divided into a training sample that consisted of 70% of the trials and a validation sample that consisted of the remaining 30%. The processing steps described below were performed separately for the training and the validation trials. To minimize random effects caused by splitting of the datasets, we performed this procedure 50 times resulting in 50 training datasets and corresponding 50 validation datasets.

It is important to note that different neurons within the sample were recorded during separate sessions and the preferred saccade directions and amplitudes could differ between the measurements dependent on the properties of the investigated neuron. Consequently, the recorded samples cannot be used to decode the full vector of the saccades but only their end-positions which were always similar between the recordings. To combine the activities of all neurons to create the model we first had to align their activity profiles to the same spatial and temporal coordinates. The procedure is illustrated in Figure 2a, b. We first calculated continuous peri-saccadic response functions for each target position in the stationary paradigm by convolving the spike times (relative to the saccade onset) with a Gaussian (σ = 20 ms), and averaging over all collected trials. Then, we interpolated the neuronal activities for x/y-positions in-between the measured saccade end positions using linear interpolation. When an extrapolation of the activities was necessary to obtain a tuning over the same spatial area for each neuron, we chose the next-neighbor extrapolation method to not overestimate the activity outside of the investigated area. The resulting data matrix for each neuron sampled the area around the center of the screen from −4° to +4° in 0.1° steps for the horizontal and the vertical dimensions and the peri-saccadic times from 400 ms before the saccade onset to 350 ms after saccade onset in steps of 10 ms.

**Figure 2:**
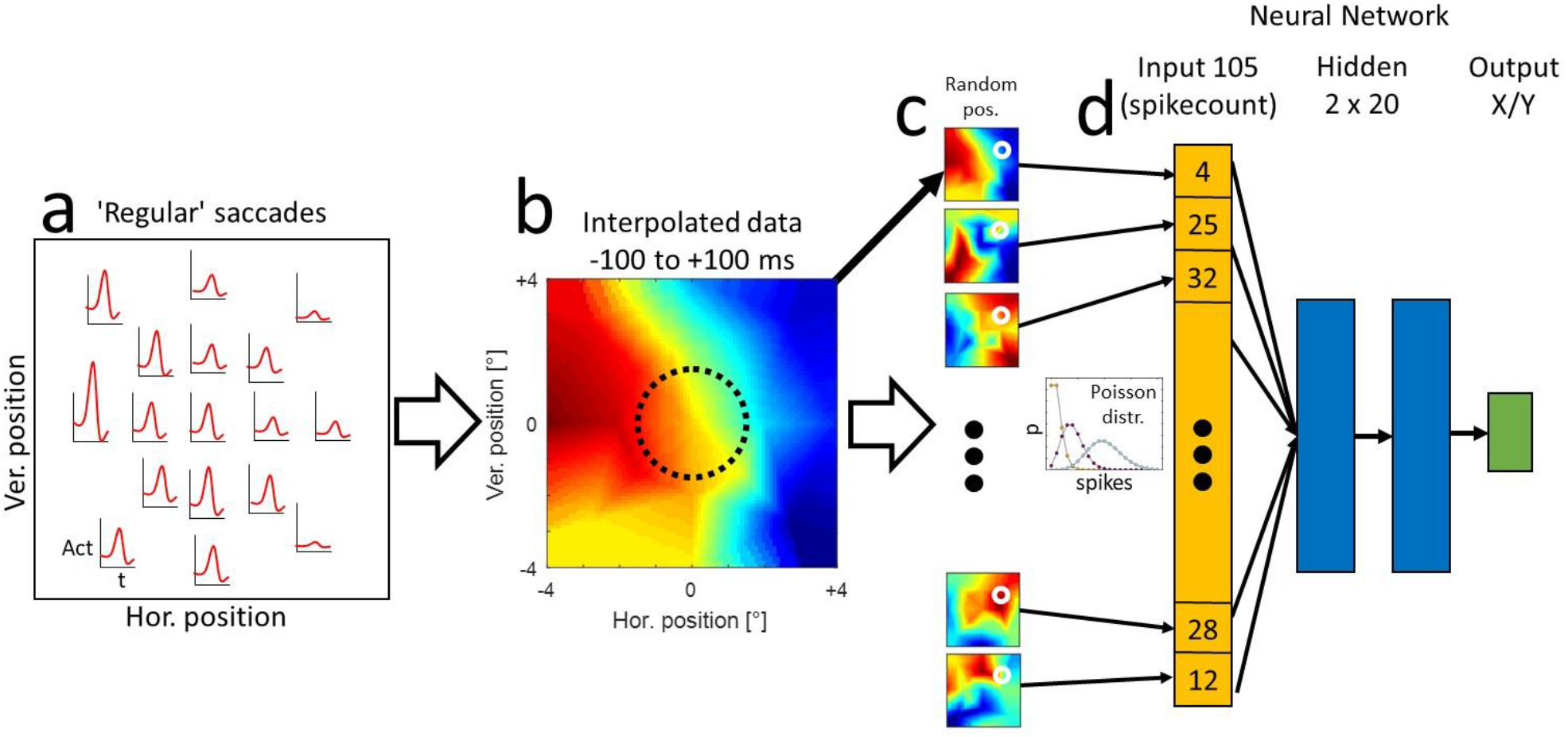
Processing steps towards a model predicting the end-position of regular saccades from activity of the sample of 105 neurons. a: For one example neuron the peri-saccadic activity was sampled from 17 positions around the center of the screen. b: The activities were then interpolated over space and time in the range of +- 4° in horizontal and vertical directions. The average activity in a time window of +- 100 ms around the saccade onset was used for the training of the neural networks. The dotted circle represents a smaller area in which the interceptive saccades and regular saccades were later compared. c: For each training dataset a random spatial position was chosen and the activity of each of the investigated neurons at that position (white circles) was used as a mean of a Poisson distribution from which the number of spikes in each trial was randomly drawn. d: The randomly generated trials were used to train the neural network model consisting of 105 elements in the input layer and 2 hidden layers with 20 elements each to predict the horizontal and vertical coordinate of the landing position of the saccade.

The same procedure was used for estimating the spatio-temporal activity profiles during the interceptive trials, using 8 saccade end positions that resulted from the different directions of target motion. Thus, the spatial area in which the interceptive saccades were investigated was smaller than for regular saccades since the saccade end-points were deflected by the stimulus motion on average only by ^~^1.5° (see Figure 3) so the relevant area was only ^~^1.5° in each direction around the center of the screen.

**Figure 3:**
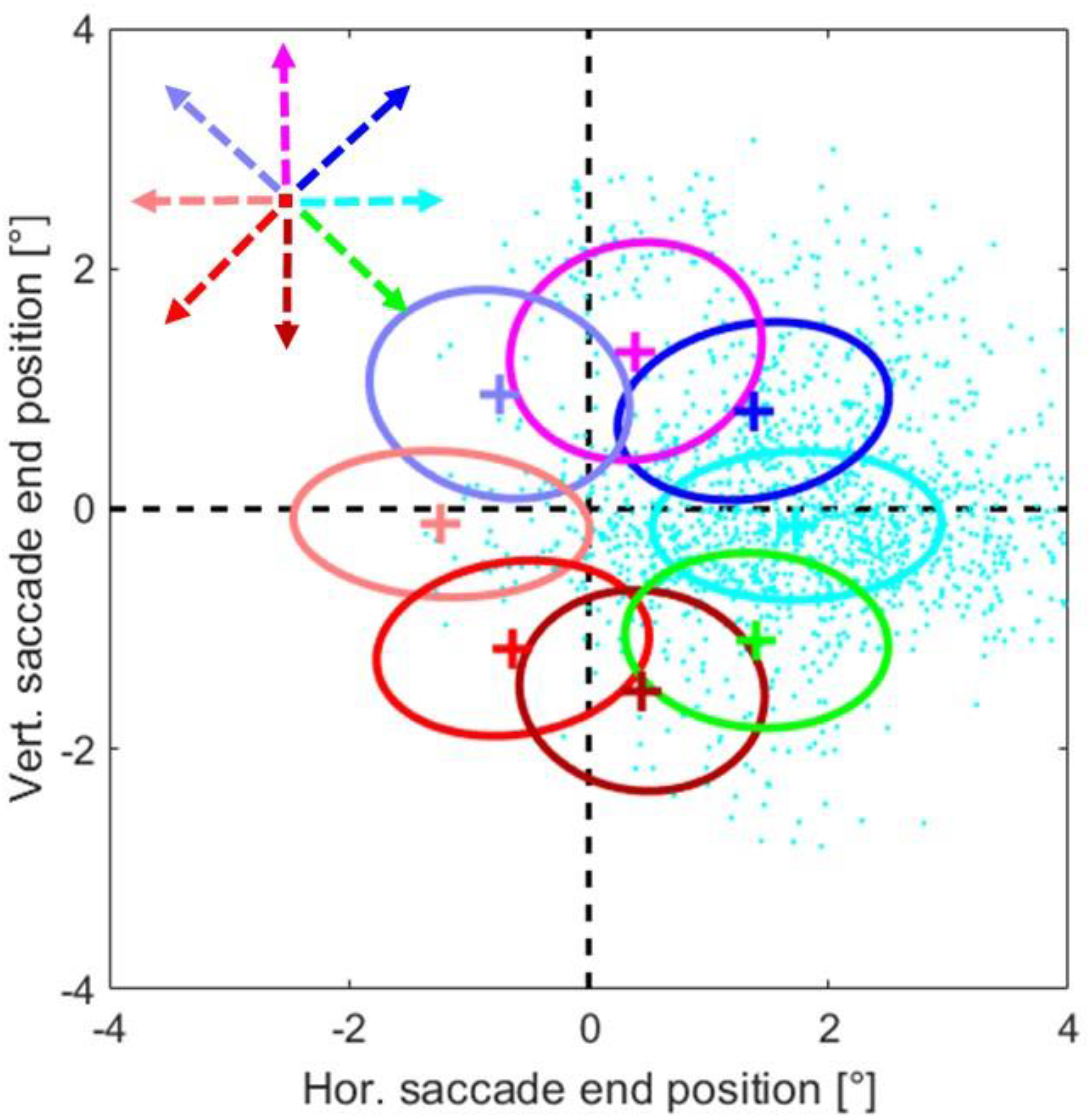
Two dimensional Gaussians fitted to the end-positions of interceptive saccades. Different colors represent different target directions, crosses show the mean saccade end position and the borders of the ellipses represent one standard deviation around the mean. Raw data depicting the end points of the saccades (cyan dots) are only shown for target movement to the right (0°).

##### Training and validation of the neural network models

For large part of the investigation we used a rather broad time-window between 100 ms before and 100 ms after the onset of the saccade - the times at which most of the saccade-related activity takes place. To investigate the time course of saccadic information we also used time-windows of 100 ms duration that were shifted in steps of 50 ms in the range between 400 ms before and 350 ms after the saccade onset. For the training of the networks a large training sample is advantageous to avoid overfitting of the data by the model. Thus we generated a large set of trials based on the estimated spatio-temporal activity profile of each neuron (Figure 2c) and the assumption that the spike counts in different trials are approximately Poisson distributed (e.g. Shadlen and Newsome, 1998). To this end we chose a random saccade end-position between −4° and +4° in horizontal and vertical directions (white circles in Figure 2c) and calculated the mean expected spike-count within the investigated time window for each neuron, based on the training sample of the recorded trials. Then we used this average spike-count as the estimate for the mean of a Poisson distribution from which the actual trial was randomly drawn. This procedure was repeated 100.000 times to sample well the whole investigated spatial area.

For a validation of the model we either generated a similar dataset based on the validation trials or, for better clarity of presentation, we investigated the model predictions at discrete saccade end locations. To allow for comparison between the predictions of the model for regular and interceptive saccades, we have chosen these discrete locations as the average end-positions of interceptive saccades for different directions of target motion (Figure 3). In this way the model was tested only at those spatial positions where sufficient data was obtained for regular as well as for interceptive saccades.

For the decoding of the saccade end points from the activities in our neuronal sample we chose a shallow neural network model as implemented in the MATLAB Neural Network toolbox (R2018a). In our model the size of the input layer was determined by the size of the neuronal sample (105) and the size of the output layer by the content of the desired output (x-position and y-position, 2 elements). For the hidden layers we have tested several configurations which did not show large differences regarding the accuracy of the predictions and chose two hidden layers with twenty elements each (Figure 2d), which have shown a good accuracy as well as a relatively fast training speed. The networks were trained using the Levenberg-Marquardt algorithm (Levenberg, 1944; Marquardt, 1963). A separate network was trained for each of the 50 splits between the training and the validation samples and for each investigated time window. After the training, the performance of the networks to predict 1) saccade end-positions of regular saccades and 2) saccade end-positions of interceptive saccades was tested. As mentioned above, for regular saccades the separately generated validation samples were used for this testing. In contrast, for the study of interceptive saccades no splitting of the data was necessary since the trials from interceptive saccades were always independent from the training trials and thus here the full data sample was used.

## Results

### Interceptive saccades

During the single unit recordings we collected the data from altogether 11631 interceptive saccades. For a brief overview - the average amplitude of these saccades was 9.7° (std 3.9°), the average latency was 134 ms (std 44 ms) and the mean duration (time between the onset of the saccade and its end) was 36 ms (std 8.9 ms). To confirm the expected effect of target motion on saccadic eye movements, we investigated the end-points of saccades towards targets moving from the center of the screen (coordinate 0°/0°) in eight different directions. We fitted 2D Gaussians to the saccade end-positions separately for each motion direction as shown in Figure 3. The results demonstrate that the motion of the stimulus exhibited a clear influence on the end-positions of the interceptive saccades. They were, on average, shifted in the direction of the target motion, however, there was a major spread of the end-positions as can be seen by the example data-points (cyan) that are provided for one motion direction. Another hallmark of interceptive saccades is the dependence of their trajectory on their timing (Quinet and Goffart, 2015; Bremmer et al., 2016), in that the target-motion dependent component becomes larger at longer saccade latencies. This dependence is confirmed in Figure 4b that shows the relationship between the time of the saccade end and the motion-related component of the saccade. For that purpose the saccade end position is expressed as the ‘Intercept’ – which is the orthogonal projection of the saccade end-position on the vector of the target-motion (Figure 4a) and was calculated as:

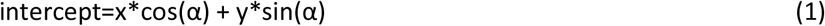

Where x and y are the coordinates of the saccade end position and α is the angle representing the direction of target motion.

**Figure 4:**
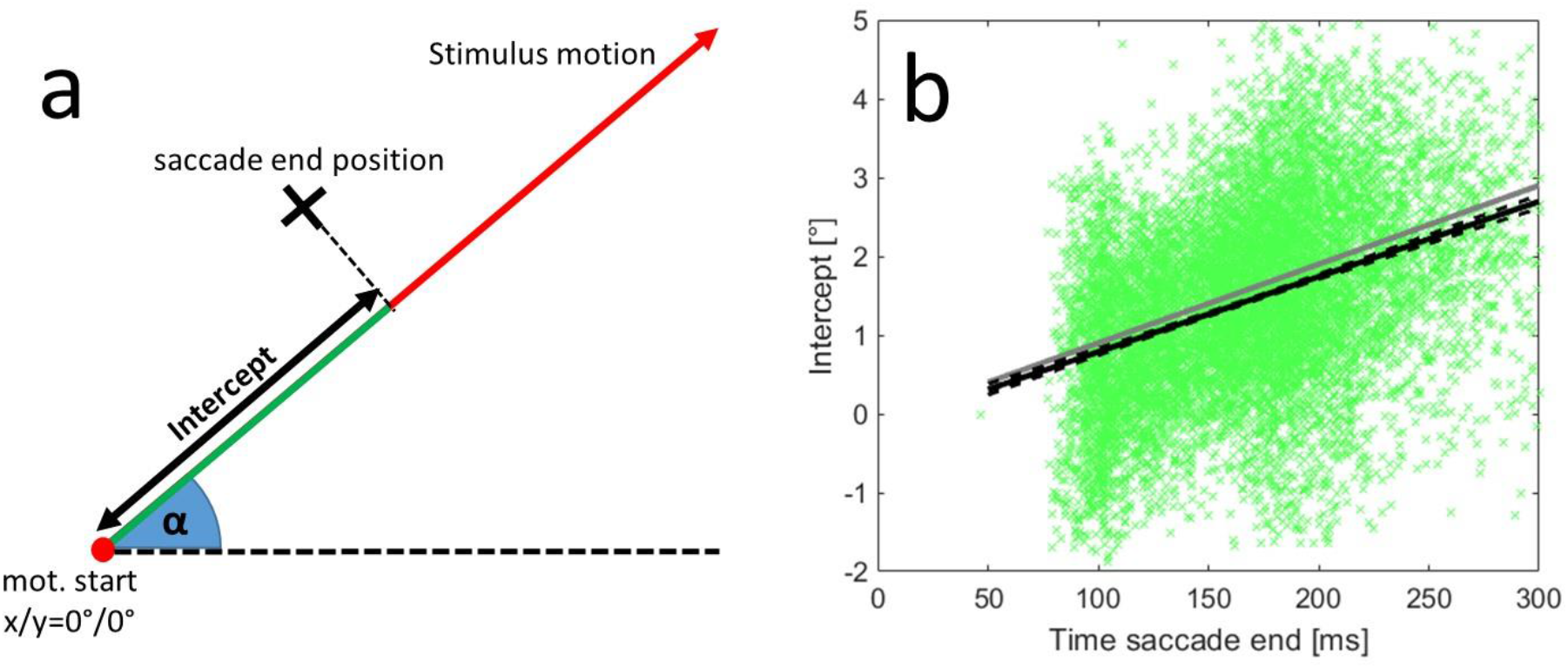
a: Description of the ‘Intercept’ as the component of saccade end-position that was aligned with the motion vector of the target. b: Relationship between the timing of the end of the interceptive saccades and their intercepts. The data from 11.631 interceptive saccades are shown as green crosses, the correlation between the time of saccade end and the intercept was 0.39 (p<0.001). The grey line marks the position of the target (relative to the center of the screen) at the respective time. The black line shows the results of a linear regression of the data and the black dashed lines its confidence interval (p=0.01, as calculated from a bootstrapping procedure). The motion directions of the targets were collapsed for this analysis, results for individual directions are shown in Figure S-1.

**Figure 5:**
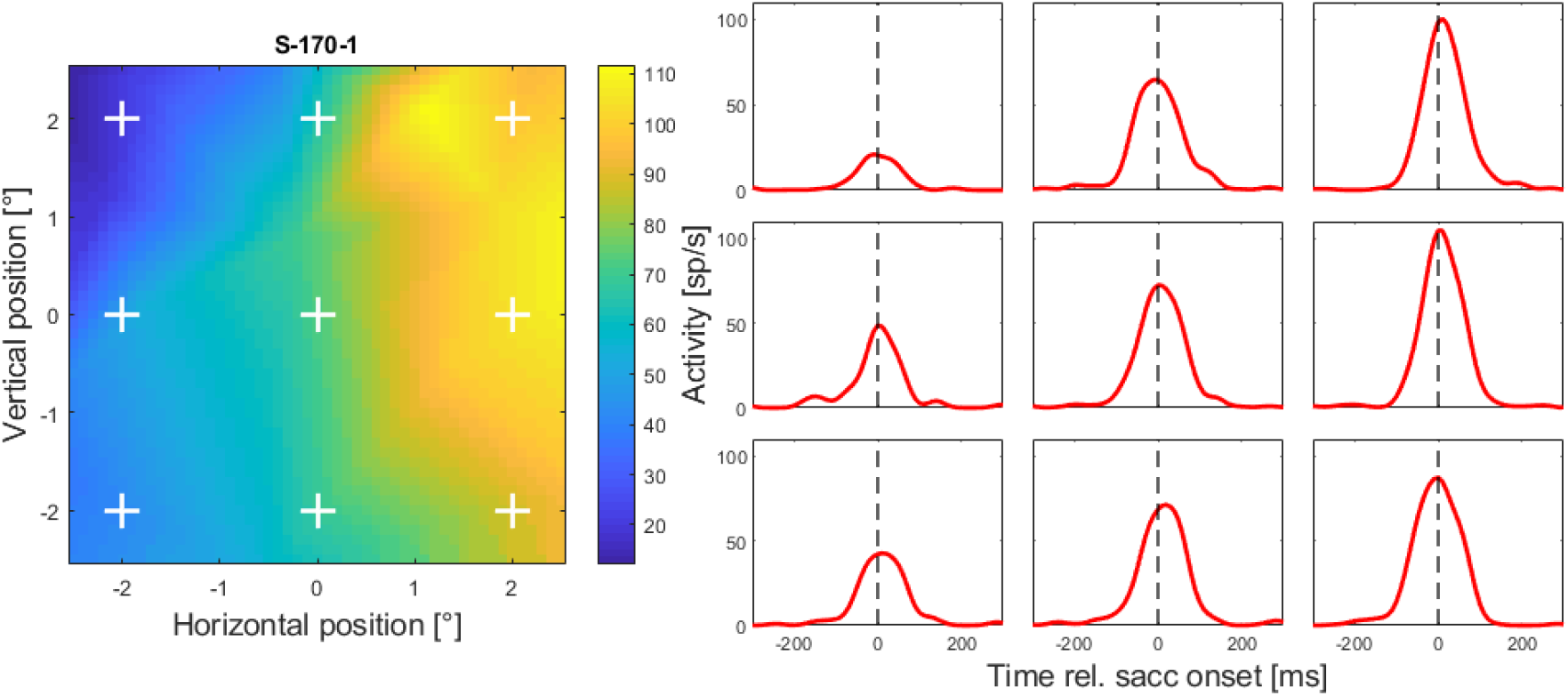
Time course of the interpolated peri-saccadic activity from one example neuron taken at nine example positions around the center of the screen. Note, that these position were chosen to give an overview over the investigated spatial area and do not coincide with the locations of the saccade targets used in the experiment. The left plot shows the spatial activity profile of the neuron at saccade onset (calculated using the spatial and temporal interpolations as described in the Methods), the white crosses marking the positions from which the temporal profile is shown in the plots on the right. The arrangement of plots on the right corresponds to the arrangement of the crosses in the left plot.

Although the landing positions in single trials have shown some variability (std of regression residuals = 0.82°), they show a clear correlation between the time of saccade end and the intercept (Pearson correlation: r=0.39, p<0.001). The regression line fitted to the data (black line) resembled the target position at the end of the saccade (grey line) which is also supported by the resulting equation of the regression in which the slope was 9.6°/s (target speed was 10°/s) and the crossing of the y-axis was at −0.17°. All target directions were collapsed for this analysis, results for individual directions are shown in Figure S-1.

### Single-unit responses

As described in the methods section, we interpolated the activities of single neurons recorded during saccades towards stationary targets as well as during interceptive saccades between different spatial positions and different times relative to the saccade onset. We focused on the area of 4×4 degrees around the center of the screen since this is the area where most of the interceptive saccades landed. The time course of the perisaccadic-activity in area LIP was in detail described elsewhere (e.g. Barash et al., 1991). Thus here we only show the temporal profiles at different spatial positions for one example neuron. The profiles show two typical features that were found in most of the neurons in the investigated sample. Firstly, the neurons reached their maximal activity around the time of saccade onset and, secondly, they show some degree of saccade-related activation at all investigated spatial locations, however, the amount of activation varied between locations.

Next, we compared the activity profiles obtained from the regular and the interceptive saccades. The activity maps at the time of the saccade onset from four example neurons are shown in Figure 6 (more examples from another four neurons are shown in Figure S-2). From a visual inspection, activity profiles from two neurons shown in the left two columns appear to show a good resemblance for regular and interceptive saccades while profiles from two other neurons shown in the right columns show no such resemblance. We used two quantitative measures to calculate the degree of similarity between the profiles. A Pearson correlation between the profiles which was 0.75 and 0.87 for the neurons in the left columns and 0.12 and −0.17 for the neurons in the right columns. We also calculated a similarity index (Sotero et al. 2010) as:

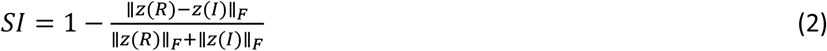

where ║X║_F_ is the Frobenius norm of X, z(R) is the z-transformed activity profile for regular saccades and s(I) is the z-transformed activity profile for interceptive saccades. The value of SI is always between 0 and 1, where 1 would be only achieved when R and I were identical. In the examples in Figure 6 the SIs of the neurons in the left column were 0.43 and 0.70 and in the right column 0.12 and 0.16. For more robust results we next averaged the activity in the time window between 100 ms before and 100 ms after the saccade onset and used these maps to compare the activity profiles for all neurons in our sample. The results are shown in Figure 7. The correlations and the SI-values point towards a continuum in degrees of similarity between the activity profiles for the two types of saccades. At least in a part of the sample of neurons saccade-related tuning appears to be very similar, independent of whether the saccade end position was systematically changed by different positions of stationary saccade targets or by interception of targets moving in different directions.

**Figure 6:**
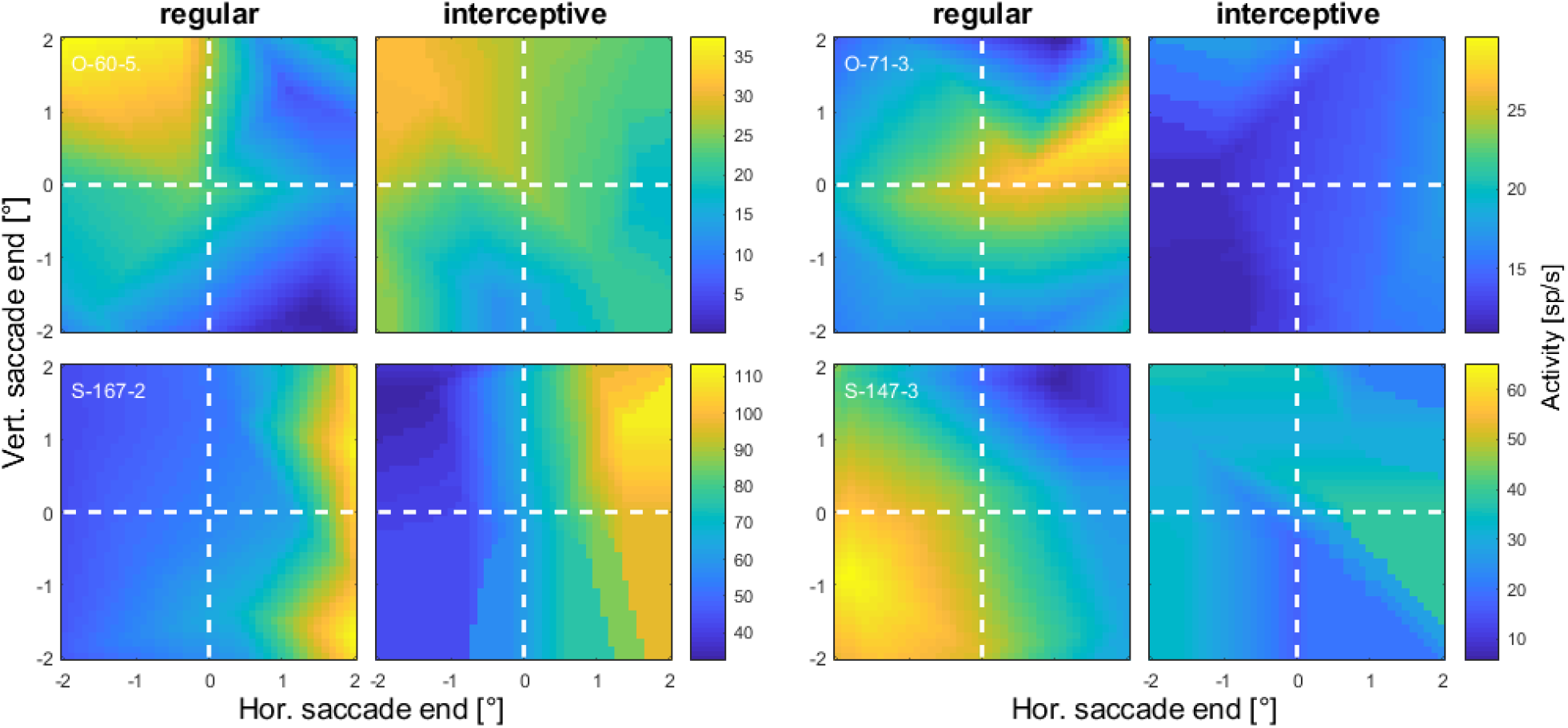
Spatial tuning profiles from four example neurons calculated from a Gaussian window (σ = 20 ms) centered on saccade onset for regular and interceptive saccades. The positions at the x and y axes represent the locations of saccade endpoints relative to the center of the screen. The two examples in the left two columns show an apparent similarity between the two activity profiles while for the neurons whose activity is shown in the two columns on the right no such similarity was observed. Note that the color scales were chosen rather to optimally compare the tuning profiles within single neurons than to make comparison between the neurons. Data from another four example neurons are shown in Figure S-2.

**Figure 7:**
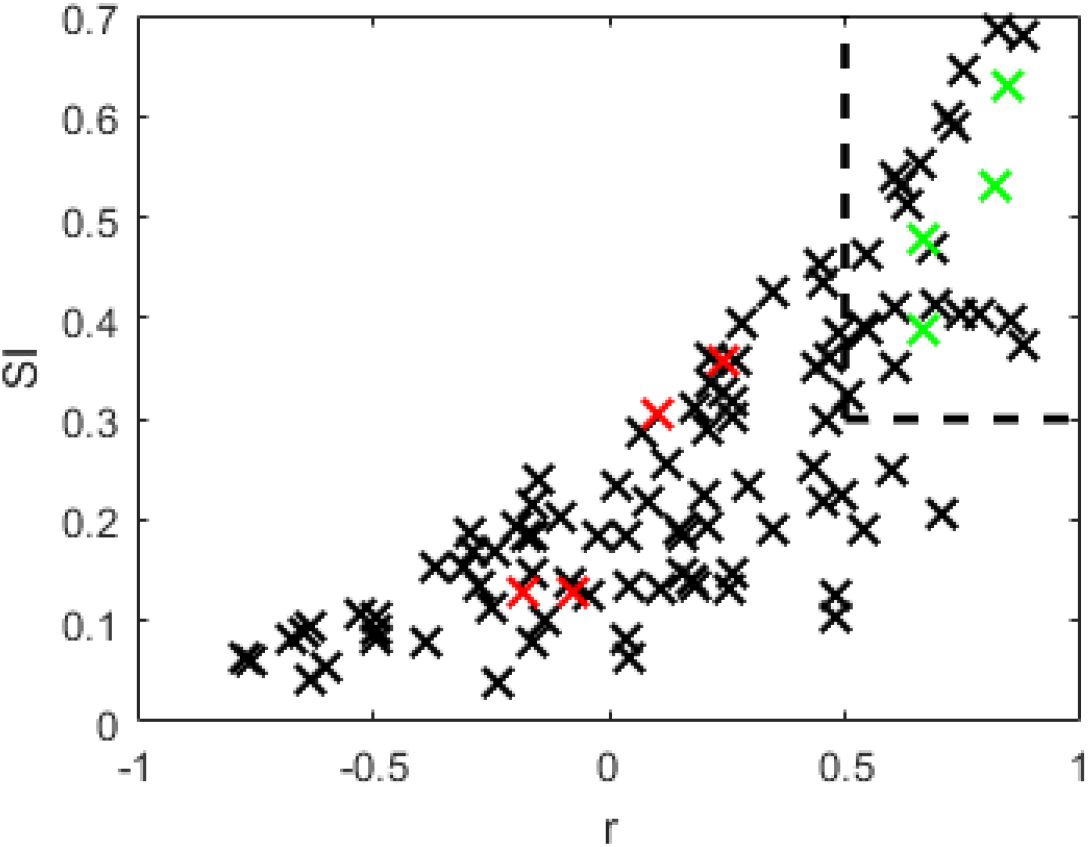
Measures of similarity between the activity profiles for saccades to stationary targets and interceptive saccades for a sample of 105 LIP neurons. The activity profiles were averaged in a time-window between 100 ms before and 100 ms after saccade onset. The Pearson correlation coefficients are plotted at the x-axis and the similarity coefficients (as defined in Eq. 1) on the y axis. The red and the green crosses mark the neurons that are shown as examples in Figures 6 and S-2. Dashed lines mark the area containing 25 neurons that were used in a later analysis of neurons with highest similarity of profiles between regular and interceptive saccades.

### Representation of regular and interceptive saccades in LIP population activity

We trained neural network models to predict the landing points of saccades towards stationary targets from the activities of our sample of 105 LIP neurons. As described in the Methods, we split the data in 70% training and 30% validation trials 50 times to mitigate the effects of random differences between the splits. Figure S-3a shows an example of a validation test of one of the resulting 50 networks that was trained on activity in the time-window between −100 and +100 ms relative to the saccade onset. As stated above, we decided to test the performance of the model at locations that represented the average landing points of interceptive saccades because here sufficient data was collected for the regular as well as for the interceptive saccades and thus a fair comparison of the two was possible. Figure S-3a shows the model results for the validation trials from one of the networks for regular saccades. The black crosses represent the predicted saccade end points for 1000 single trials that were generated from data at positions marked by the red circles. Means and standard deviations in x and y directions are shown as green crosses. The averages of the estimates apparently fit the tested positions well in most cases. This is also supported by the confusion matrix on the right side of the plot in which the trials are assigned to one of the eight tested positions based on the minimal Euclidian distance. On average 47% of single trials were assigned correctly (12.5% are expected by chance) and on average 88% were assigned correctly or to one of the direct neighbors (37.5% are expected by chance). Some of the tested positions (e.g. position 3) show a systematic error of the predictions which likely is a result of random effects of splitting the original data into training and validation samples. In Figure S-3b, the same neural network as in S-3a was tested using data from interceptive saccades. Also here the distributions of predicted saccade end positions deviate from the center towards the tested positions. This is also confirmed by the confusion matrix showing on average 36% of correct assignments and 75% of assignments that were either correct or one of the direct neighbors. In this case, however, a clear bias towards the center of the screen can be observed for all of the tested positions.

We combined the predictions for all 50 networks that were trained using the different data splits. The results are shown in Figure 8. Here the mean predicted positions from all tested networks are marked as a cross and the averaged standard deviations for all tested networks in x and y directions are shown as an ellipse. As in Figure S-3, we found that the predicted end positions of regular saccades (Figure 8a) were on average quite accurate with a mean error of 0.38° (for comparison - the typical accuracy of the EyeLink eye-tracker is reported to be 0.25° to 0.5°). The confusion matrix showed on average 37% correct assignments and 81% correct or direct neighbor assignments. We also found that the neural networks were capable to predict saccade end positions of interceptive saccades (Figure 8b); here, the confusion matrix showed 34% of correct assignments and 74% correct or direct neighbor assignments. However, a systematic error towards the center was also observed for all tested positions. For an easier comparison between the two types of saccades we calculated the ‘center bias’ of the predictions as the difference between the intercept (Figure 4a, eq. 1) of the tested position and the intercept of the prediction. A positive value indicates that the predicted saccade position is shifted towards the center relative to the tested position and vice versa. In Figure 8c, d the center bias is shown for the different tested positions and as averages over all positions. The center bias for interceptive saccades was on average 0.75° and significantly (p<0.01, paired t-test) higher than for regular saccades (average 0.25°).

**Figure 8.**
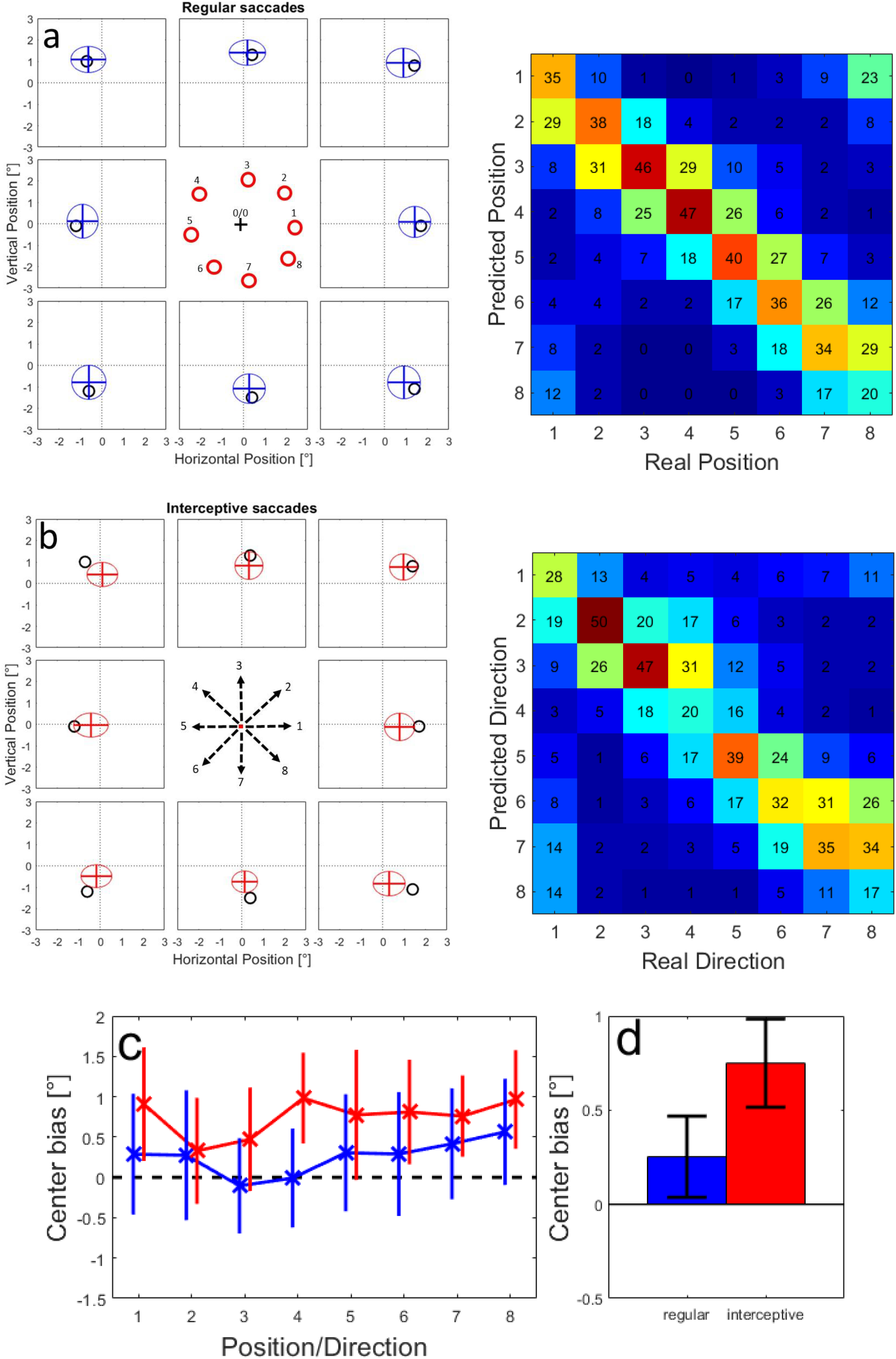
a: Validation of the training success for 50 networks that were trained on 70% of the available data from regular saccades and tested using the remaining 30%. Black circles show the tested positions, the ellipses represent the average positions and average standard deviations from 50 validation samples. In the confusion matrix on the right, the predictions from single trials over all validation samples were assigned to one of the tested positions based on the minimal Euclidian distance between the prediction and each of the tested positions. b: The same plot for predicting the end positions of interceptive saccades from the population activity based on neural network models trained using data from regular saccades. c: Averages and standard deviation of the Center bias for different tested positions for regular (blue line) and interceptive (red line) saccades. The lines are slightly shifted for better visibility. d: Averages and standard deviations of the center bias for regular (blue) and interceptive (red) saccades calculated over all tested positions.

We further investigated this systematic pattern of prediction errors by making the model networks predict saccade end-points in a continuous area in a range of 1.5° around the center of the screen (coordinate 0/0). Figure 9 (a, b) shows the error vectors between the tested position (arrow backs) and the prediction of the model (arrow tips) for regular (a) and interceptive (b) saccades. Note that in this figure, for a better overview, we have chosen to reduce the number of presented locations and reduced the length of the arrows (representing the size of the error) by 50%. The figures show that while there is no clear pattern of errors for the regular saccades, the predictions for the interceptive saccades are clearly biased towards the center. This center bias increases with increasing eccentricity of the tested positions. The details about the direction and the amplitude of the errors for all tested positions are then shown in Figure 9c-f for all tested positions. They confirm that while for the regular saccades only small (albeit systematic) errors were observed (c, e), for the errors for interceptive saccades a pinwheel structure of error directions (d) indicates a compression of the predictions towards the center of the screen. The strength of the error grows with increasing eccentricity with the exception of an area in the upper right quadrant in which the predictions were more accurate than in the rest of the tested positions (f).

**Figure 9:**
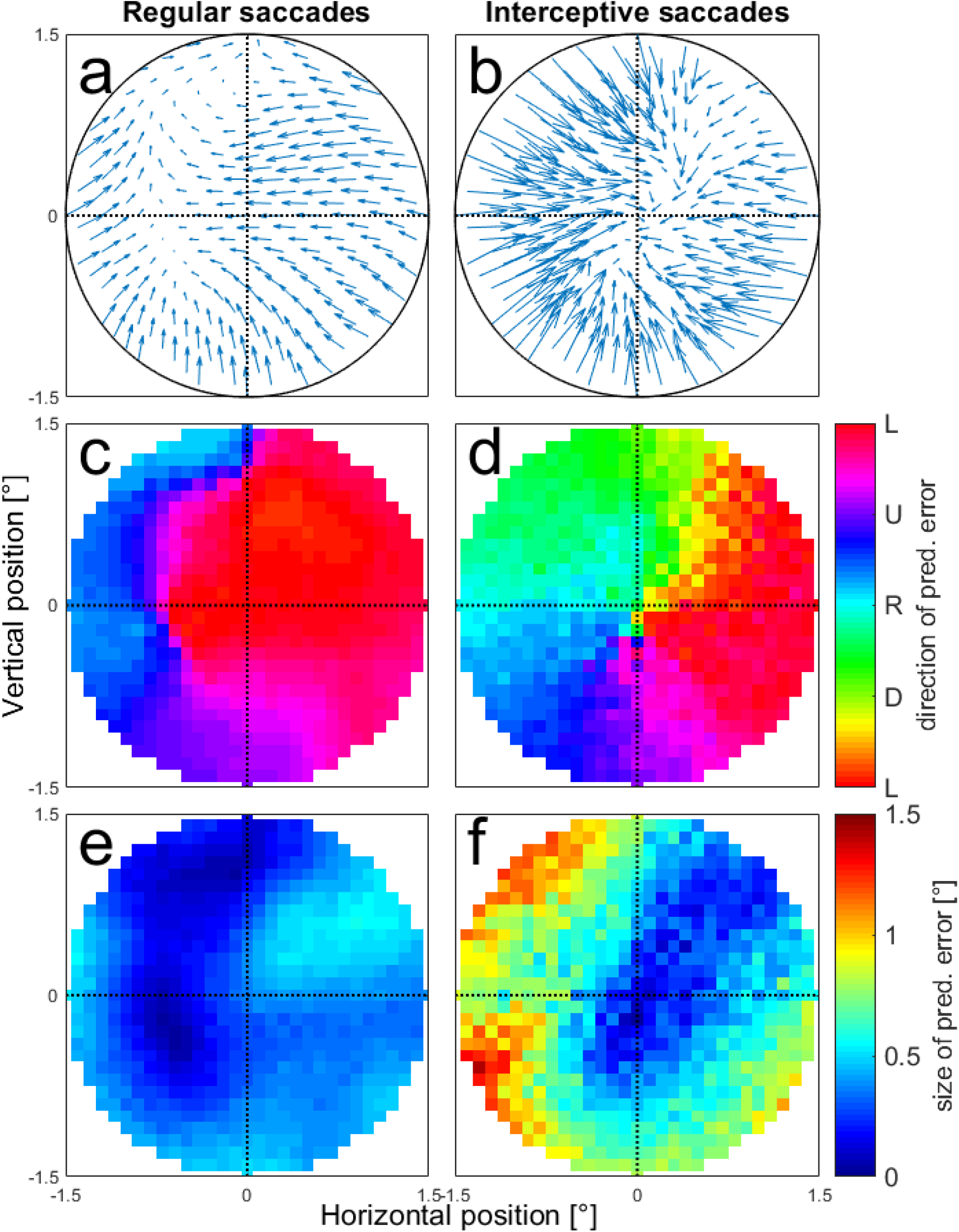
Prediction errors of saccade end positions tested at different positions in the area of 1.5° around the screen center. a, b: Average error vectors shown at selected positions. The arrows indicate the differences between the tested positions and the predictions of the model. The length of the error vectors is scaled down by 50% for a better overview. c, d: Average directions of the error vectors for regular (c) and interceptive (d) saccades. e, f: Average length of the error vectors for regular (e) and interceptive (f) saccades.

Results shown in Figure 7 indicate, that there is a continuum of similarities between the activity profiles between regular and interceptive saccades in the LIP neuronal population, ranging from very different to close to identical. Thus, we asked whether these similarities also reflect the ability of the neurons to code for the end position of interceptive saccades (and consequently represent the movement of the target). To test this, we focused on a sub-sample of neurons with the strongest similarity between the two activity profiles as obtained from the regular and interceptive saccades. For this sample of ‘best’ neurons we selected 25 neurons in which the Pearson correlation between the spatial activity profiles for regular and interceptive saccades was larger than 0.5 and the Similarity index was larger than 0.3. These borders are marked by a dashed line in Figure 7. Using this sample of neurons, we again trained the neural networks for predicting the endpoints of interceptive saccades based on training data taken from regular saccades. The results are shown in Figure 10 in the same format as Figure 8. It indicates that while the predictions were on average accurate for regular saccades, the center bias that was previously observed for interceptive saccades was not fully eliminated albeit strongly diminished and was now on average only 0.3°. Given the target speed of 10°/s this spatial bias can be transformed into a 30 ms time-lag on the target, which corresponds well with the position of the target at the beginning of the interceptive saccade (average duration of interceptive saccades was 36 ms).

**Figure 10:**
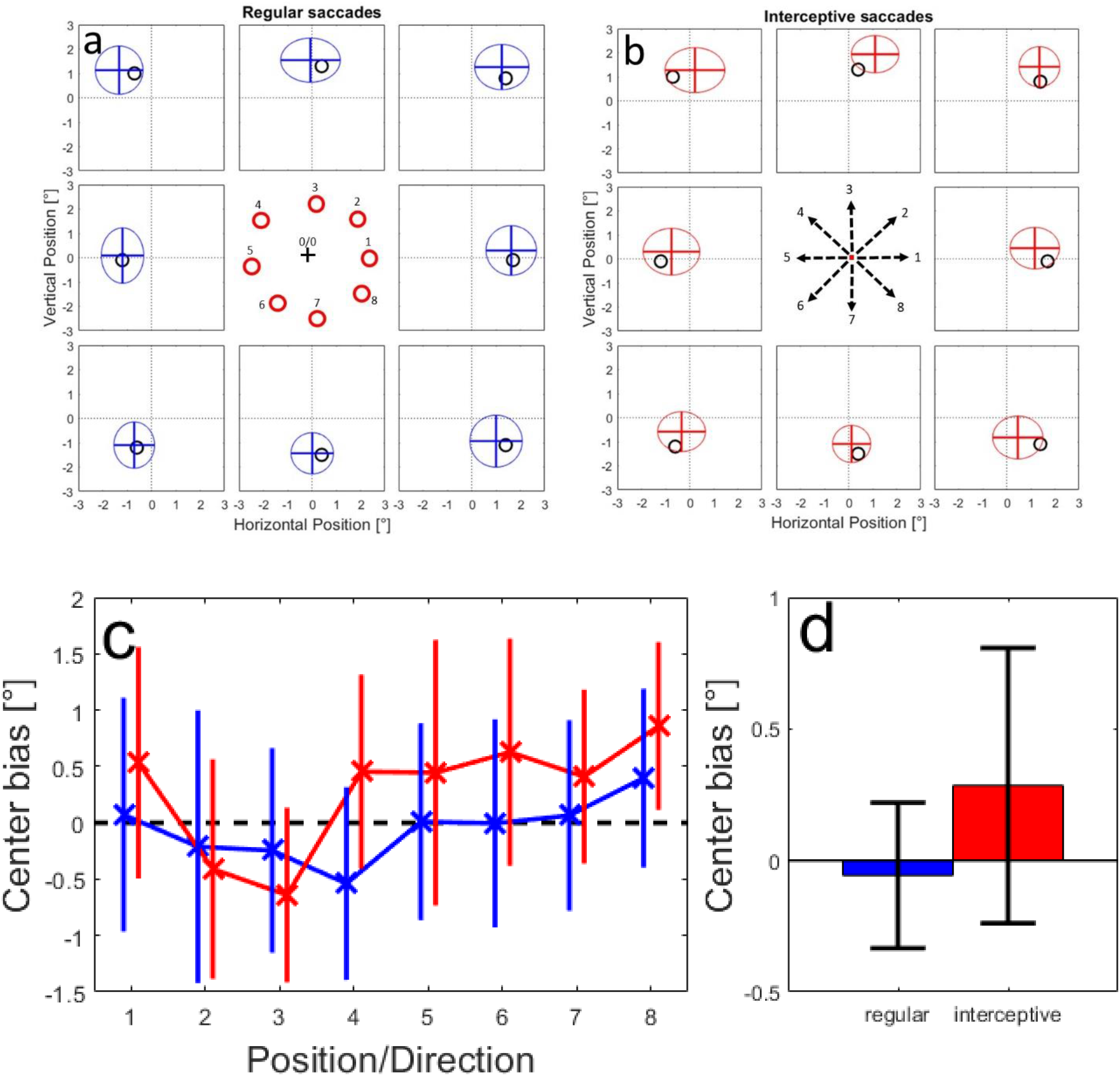
Same predictions of saccade end positions as in Figure 8. Here the networks were trained based on activity of 25 neurons with the strongest similarity between the activity profiles for regular and interceptive saccades.

### Time course of motion information processing

Up to this point we investigated the activity collected in a rather broad time-window between 100 ms before and 100 ms after the saccade onset. To obtain a temporal profile of the saccade-related information in our neuronal sample, we also trained neural networks based on activity in time-windows of 100 ms duration that were moved in steps of 50 ms starting from 400 ms before and ending at 350 ms after the saccade onset. As described in the Methods, the data in each time-window was again split fifty times into training and validation datasets and the networks were trained on the training datasets of each of these splits separately. We used the same example saccade end-positions as in previous sections to evaluate the predictions of the model for regular and interceptive saccades. The supplemental videos S-5 and S-6 show the development of the means and the standard deviations of the predictions over time in the same format as in Figure 8. The line intersections represent the average predicted saccade end-position for regular (video S-5) and interceptive (video S-6) saccades. The ellipses indicate the average standard deviations of the predictions in horizontal and vertical directions. Two effects can be observed: 1) In a narrow time-window around the onset of the saccade, the average predicted positions move from the center in the direction of the tested positions. 2) The variability of the predictions (indicated by the radii of the ellipses) decreases in approximately the same time-window. The two effects are summarized in Figure 11a, b for regular (blue) and interceptive (red) saccades. For regular saccades it shows a decrease of the center bias starting at −100 ms and reaching a minimum at saccade onset. For interceptive saccades the temporal profile appears shifted by ^~^50 ms, the center bias starts decreasing 50 ms before and reaches a minimum 50 ms after the saccade onset. Consequently the largest differences between the predictions for regular and interceptive saccades were observed before and at saccade onset. The time windows in which the center bias for the two saccade types was significantly different (paired t-test, p<0.01) are marked by asterisks. After the saccade, center bias increases again for both types of saccades as the predictions move back towards the center. No significant differences were observed in the post-saccadic time period. When investigating the trial-by-trial variability of the predictions (Figure 7b), an increase of precision (as marked by a decrease in the variability of the predictions) can be observed around saccade onset. No significant differences between the saccade types were found except for a brief period 200 ms after saccade onset to which we do not assign any specific meaning. Next, we used the same procedure on the subpopulation of 25 neurons that have shown the highest similarity of tuning profiles between regular and interceptive saccades as described above. The results are shown in supplemental videos S-7 and S-8 and summarized in Figure 11c, d. For the center bias (Figure 11c) similar differences between the temporal profiles for the two saccade types are shown as in the whole sample of neurons (Figure 11a). Also here, the pre-saccadic coding of the saccade end position for interceptive saccades is delayed by ^~^50 ms relative to regular saccades but now having the smallest bias at the same time, i.e. saccade onset. While there was a substantial center bias, for the predictions of interceptive saccades based on the whole neuronal sample, this bias was close to zero at the saccade onset when the model was only based on the selected sample of 25 neurons. The temporal profile of the variability of the predictions (Figure 11d) was less pronounced and has shown a larger post-saccadic variability for the predictions of interceptive saccades. This might be caused by the behavioral variability induced by the smooth pursuit eye movements (including catch-up saccades) that followed the interceptive saccade in contrast to a steady fixation that was following the regular saccades.

**Figure 11.**
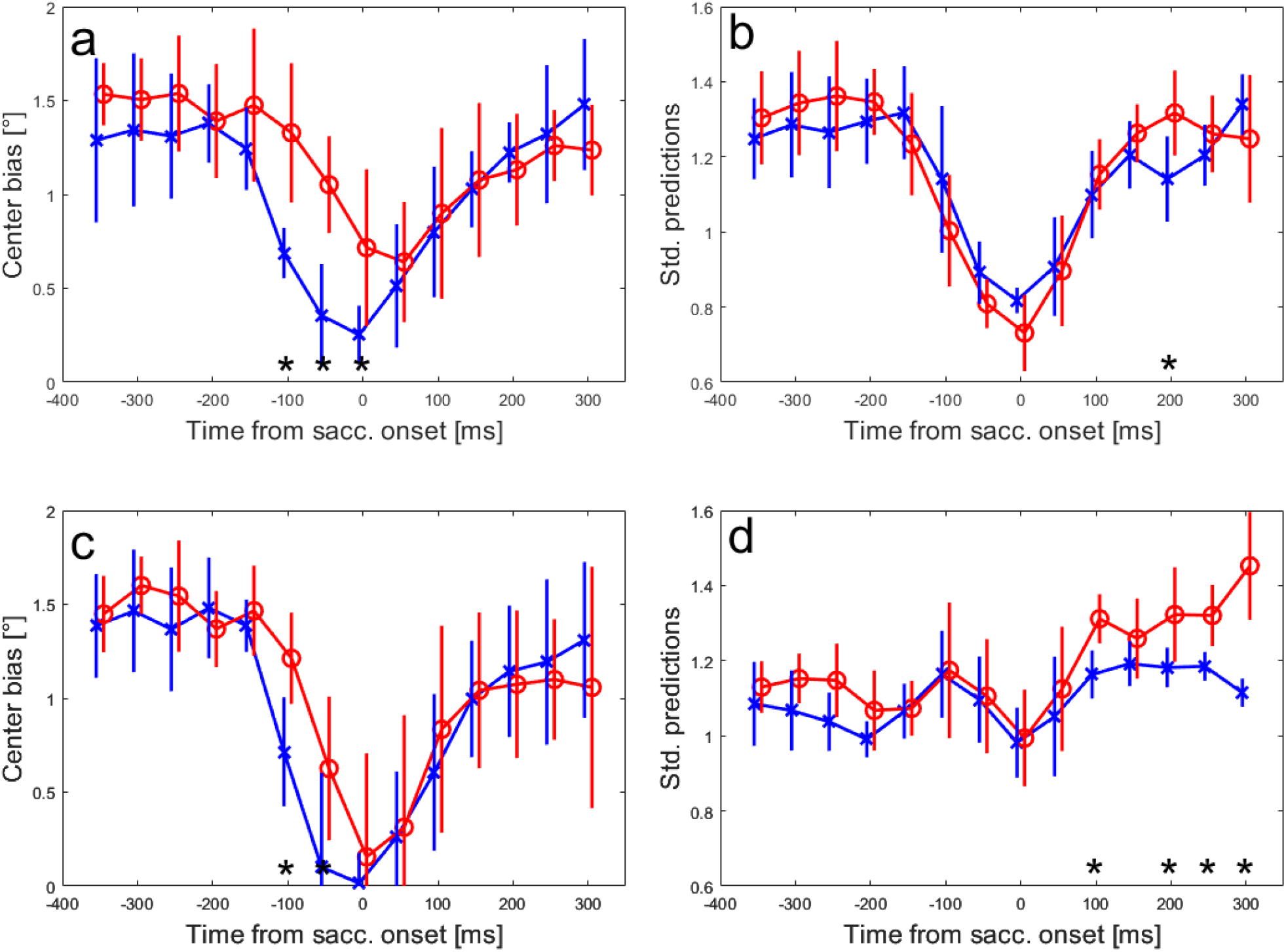
a: Average center bias of predicted end-points for regular (blue) and interceptive (red) saccades in different time-windows relative to saccade onset. The x-axes shows the center of a 100 ms time-window from which the results were calculated, the error bars show standard deviations over the eight tested positions. b: Average standard deviations of the predicted saccade end positions for regular (blue) and interceptive (red) saccades. c, d: Show the same measures as a, b, only based on the predictions derived from a sub-sample of 25 neurons that have shown the strongest similarity between the activity profiles of regular and interceptive saccades. The asterisks at the bottom of each plot indicate time windows with significant (paired t-test, p<0.01) differences between the two saccade types.

## Discussion

### Role of area LIP in generation of interceptive saccades

Our results together with our earlier study (Bremmer et al., 2016) clearly point towards an involvement of LIP neurons in the representation of interceptive saccades. Our results are clearly inconsistent with the notion that LIP neurons only code for a ‘snapshot’ of the target position taken just after the initial target step since the motion direction of the target after this step can be successfully recovered from the neuronal activity. Given this principal finding it should be, however, more precisely specified to what degree LIP neurons contribute to the very accurate interception of the target position at the end of the saccade. It is, e.g., possible, that although the neurons do not represent a snapshot of the target position taken immediately after the target step, they still may use a snapshot taken at some later point in time, when the target already moved for a distance in the given direction, and thus partly represent the motion of the target without explicitly processing the motion signal. We believe that this interpretation of our results is unlikely. In our sample of neurons we found a continuum regarding the representation of the stimulus motion by single neurons. When we predicted the saccade landing position of interceptive saccades based on all investigated neurons, the predicted landing position lagged on average by 0.8° which corresponds to 80 ms of target motion, which is consistent with a position snapshot taken ^~^50 ms before the saccade onset. However, when only those ^~^25% of neurons with the strongest similarity between the tuning profiles for regular and interceptive saccades, the time was reduced to ^~^30 ms which is approximately the time of saccade onset. It is not clear whether, given an even more extensive sampling of LIP neurons, another sub-population would have emerged, that would accurately represent the target position at the time of saccade end – and thus the actual landing position of an interceptive saccade. The current sample allows for coding of saccades that represent the target position up to the saccade onset, which gives some credibility to the idea that (given the processing latencies from LIP to the motor neurons) the neurons did not operate based just on a snapshot but on a genuine processing of a motion signal (Assad and Maunsell, 1995).

Our findings resemble an earlier report from the intermediate and deep layers of the SC (Goffart et al., 2017) that has also shown a continuum in degrees to which neurons changed the position of their motion fields during interceptive saccades. The SC is by far the most investigated area regarding its contributions to the generation of interceptive saccades. An early report (Keller et al., 1996) has shown that the preferred motion vector of saccade-related SC neurons moved in the direction of target motion during interceptive saccades relative to saccades to stationary targets. This finding can be interpreted as a lack of contribution of SC-neurons to the accurate metrics of interceptive saccades and thus a second drive was postulated that contributes the target-motion related component of the saccade.

Areas LIP and SC are strongly interconnected (Andersen et al., 1985) so that it is possible that the properties reported by Goffart et al. are inherited from area LIP. Both reports suggest that a second drive that further increases the accuracy of interceptive saccades is likely. It is not clear so far, whether this second drive comes from the activity of the Fastigial nucleus of the cerebellum as it was hypothesized earlier (Optican and Quaia, 2002; Optican and Pretegiani, 2017) or whether it is contributed by the FEF, where motion information was reported (Barborica and Ferrera, 2003). These possibilities are of course not exclusive.

### Different approaches for decoding of saccades from neuronal activity

In this manuscript we present a population-centered approach to the coding of interceptive saccades. This is slightly different from previous works (e.g. Keller et al. 1996, Goffart et al. 2017) that focused on the accurate description of single neurons their discharging properties and motion fields.

We acknowledge, that other models can be used to decode saccade trajectories from the activity of a neuronal sample. In previous publications (Bremmer et al. 2016; Churan et al. 2019) we used a maximum likelihood approach for the same purpose. In that approach we generated probability maps of different saccade end-positions based on the activity of each neuron and generated a population prediction by combining all probability maps for the specific activity patterns that were found around the time of the interceptive saccades. In particular, in Churan et al. (2019) this approach was used on partly the same data as in this manuscript. Despite the large methodological differences, the basic conclusions derived from these two approaches were very similar. Our current approach, however, allowed more clear cut results that appeared more stable (e.g. across different neuronal samples) and independent of outliers than they were when the maximum likelihood approach was used.

### Future directions

As mentioned above, our findings point in the same direction as previous results from the superior colliculus (Goffart et al., 2017). The methods chosen in that study were different from ours and thus it would be intriguing to use our decoding approach on a sample of SC-neurons to investigate whether the coding accuracy for regular and interceptive saccades is similar in SC.

A recent development has shown a promising way to further disentangle the contributions of target position and target motion for generation of interceptive saccades. Goettker et al. (2019) have demonstrated that when human subjects were asked to perform interceptive saccades towards isoluminant targets, the interceptive saccades become inaccurate and lag behind the moving target by ^~^100 ms. This is attributed to a lack of motion information provided by isoluminant stimuli (Cavanagh et al., 1984; Lu et al., 1999) to areas of the dorsal processing stream like the middle temporal, and middle superior temporal areas (Thiele et al., 2001; Riecansky et al., 2005). Thus it would be intriguing to investigate how the processing of interceptive saccades in LIP and SC changes between luminance defined and isoluminant stimuli.

## Declarations

### Availability of data and code

The datasets generated during and/or analysed during the current study as well as the code used for evaluating the data are available from the corresponding author on reasonable request.

### Ethics approval

All procedures had been approved by the regional authorities (TVA Nr.: V54-19 c 20 15 h 01 MR 13/1 Nr. G71/2017) and were in accordance with the published guidelines on the use of animals in research (European Communities Council Directive 2010/63/EU).

## Acknowledgements

This work was supported by Deutsche Forschungsgemeinschaft (CRC/TRR-135/A1 [project number 222641018], IRTG-1901, and RU 1847/A2) and by the Hessisches Ministerium für Wissenschaft und Kunst (HMWK; project ‘The Adaptive Mind’).

## Supplement

**Figure S-1:**
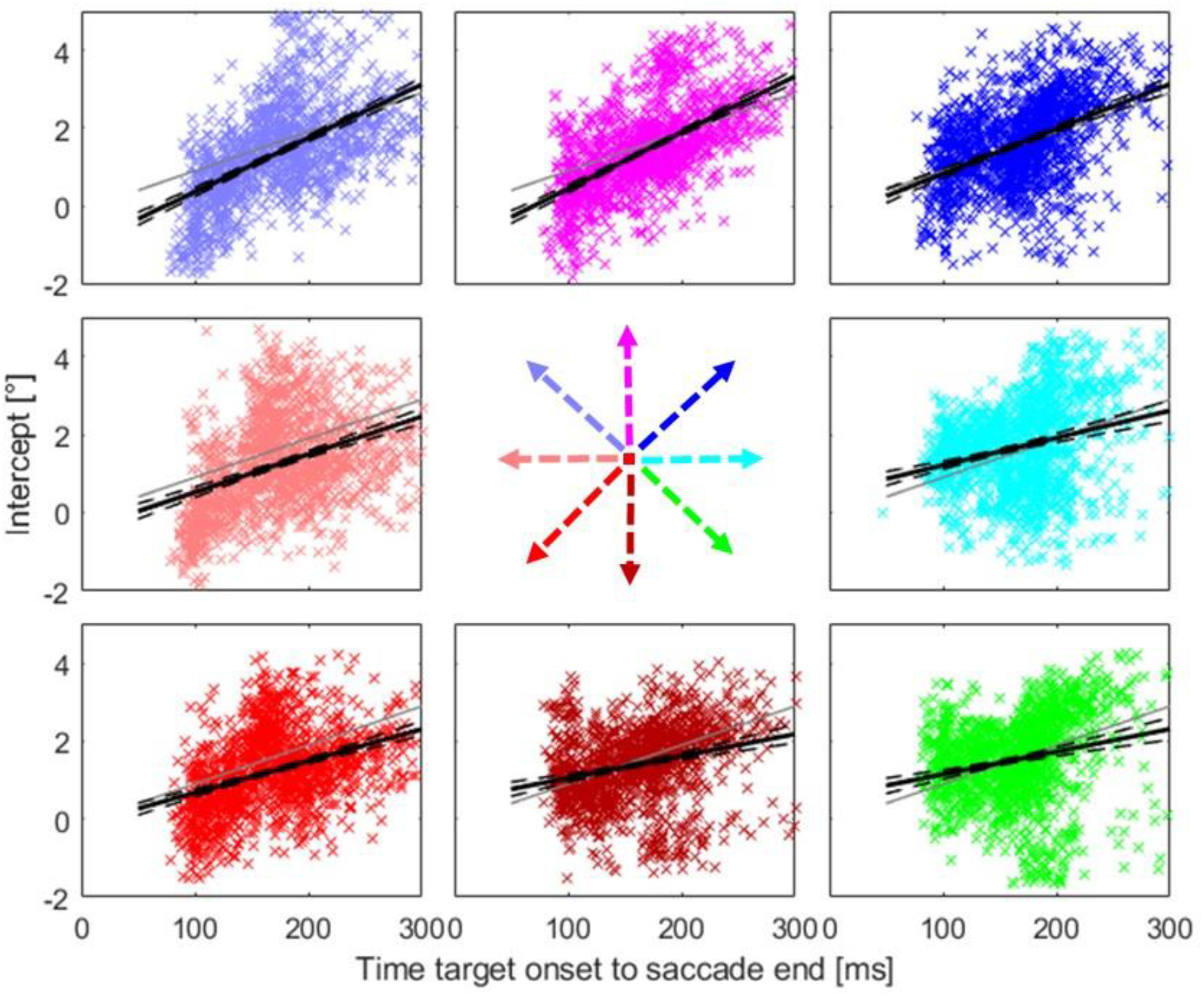
Relationship between the timing of the end of the interceptive saccades and their intercepts for different directions of target motion. Different plots show the data from individual target directions according to the arrows. Grey lines marks the position of the target (relative to the center of the screen) at the respective time. Black line shows the results of a linear regression of the data and the black dashed lines its confidence interval (p=0.01, as calculated from a bootstrapping procedure).

**Figure S-2:**
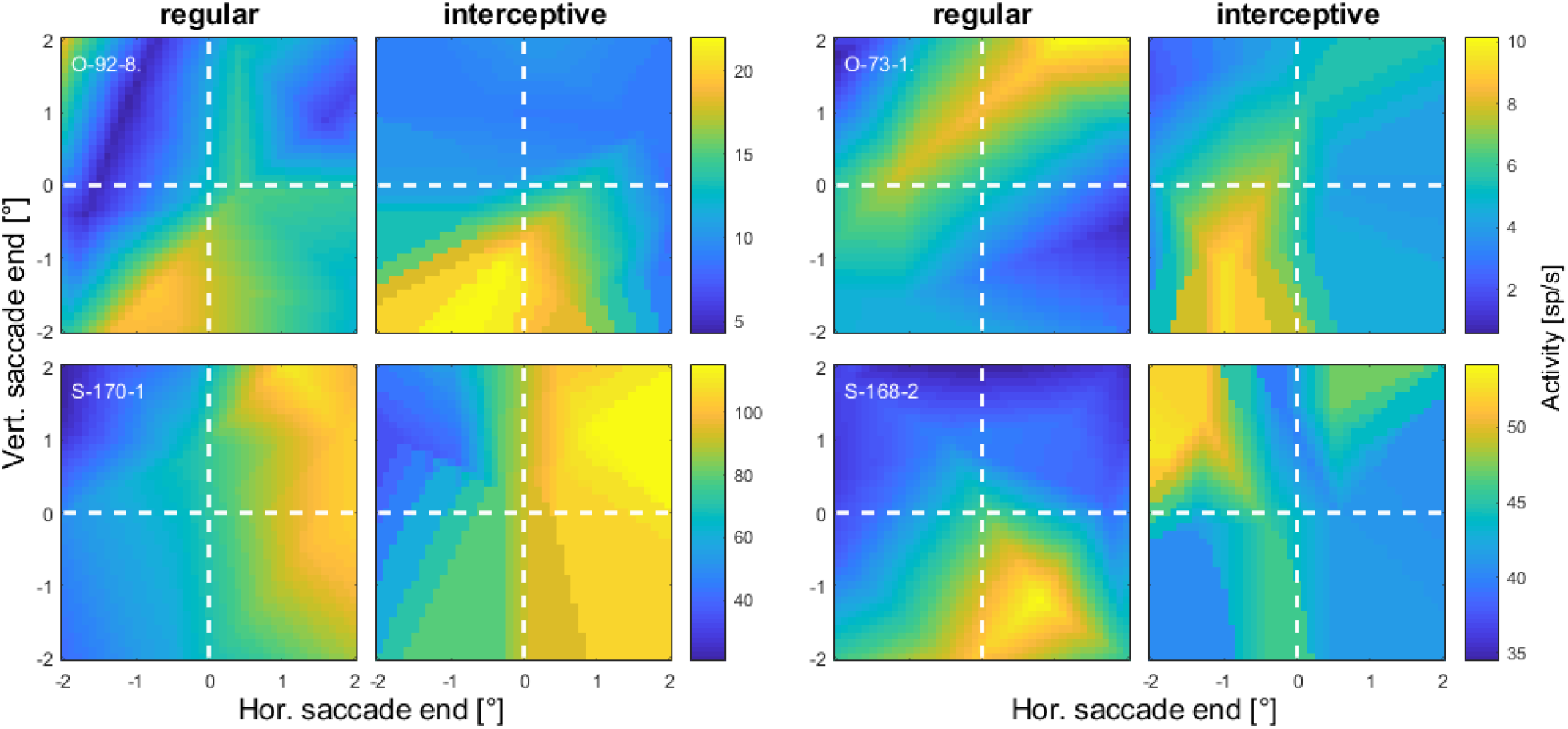
Spatial tuning profiles from another four example neurons in the same format as Figure 6. The positions at the x and y axes represent the locations of saccade endpoints relative to the center of the screen. The two examples in the left two columns show an apparent similarity between the two activity profiles while for the neurons in the two columns on the right no such similarity was observed.

**Figure S-3.**
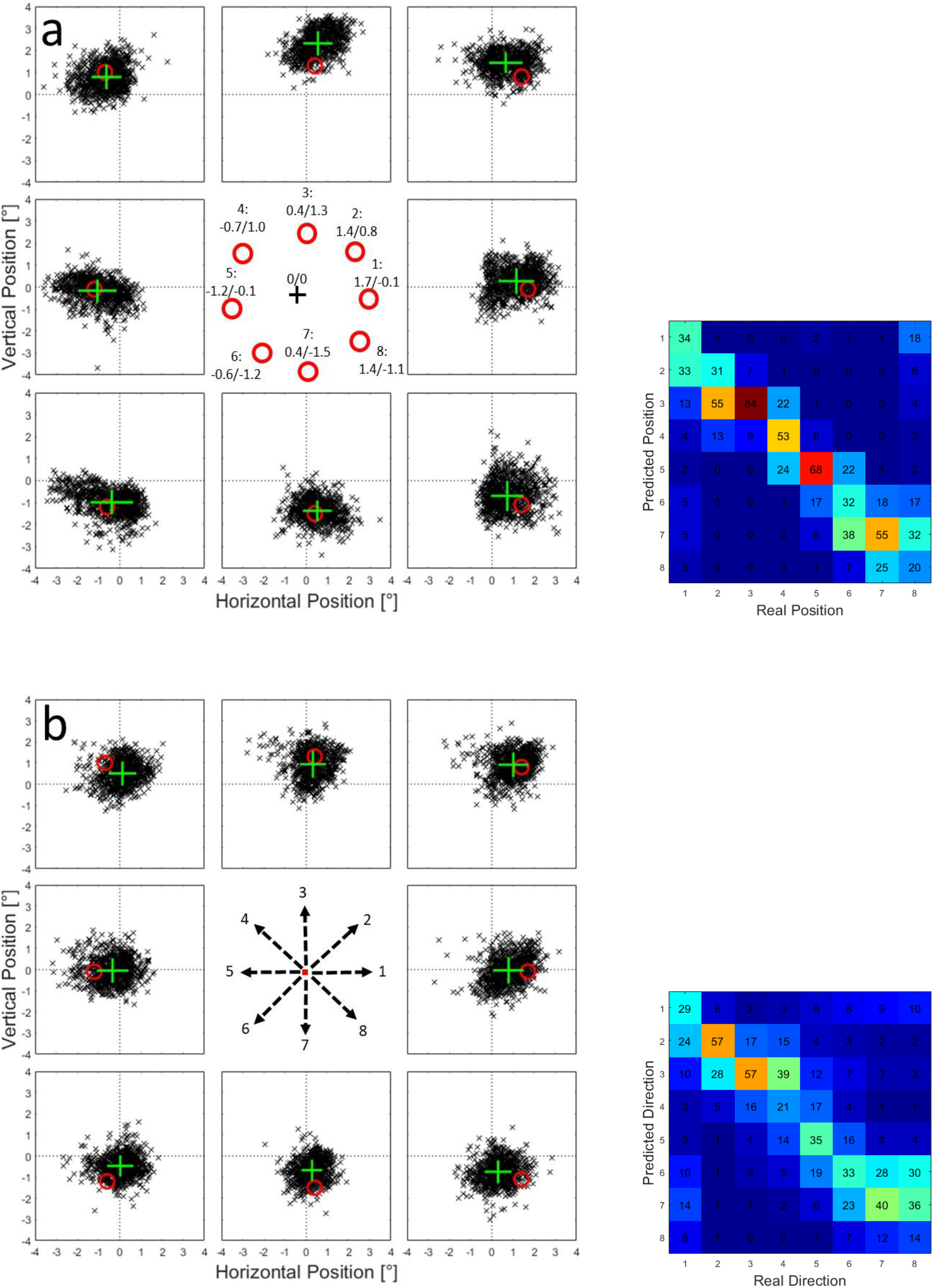
a: Example of the predictions from one neural network using estimated spike counts in a time window between 100 ms before and 100 ms after the onset of regular saccades. Black crosses represent the predictions of 1000 trials that were generated from the activity estimates for each tested position. b: Predictions of same network as in a) for saccade end-points of interceptive saccades.

**Figure S-4:**
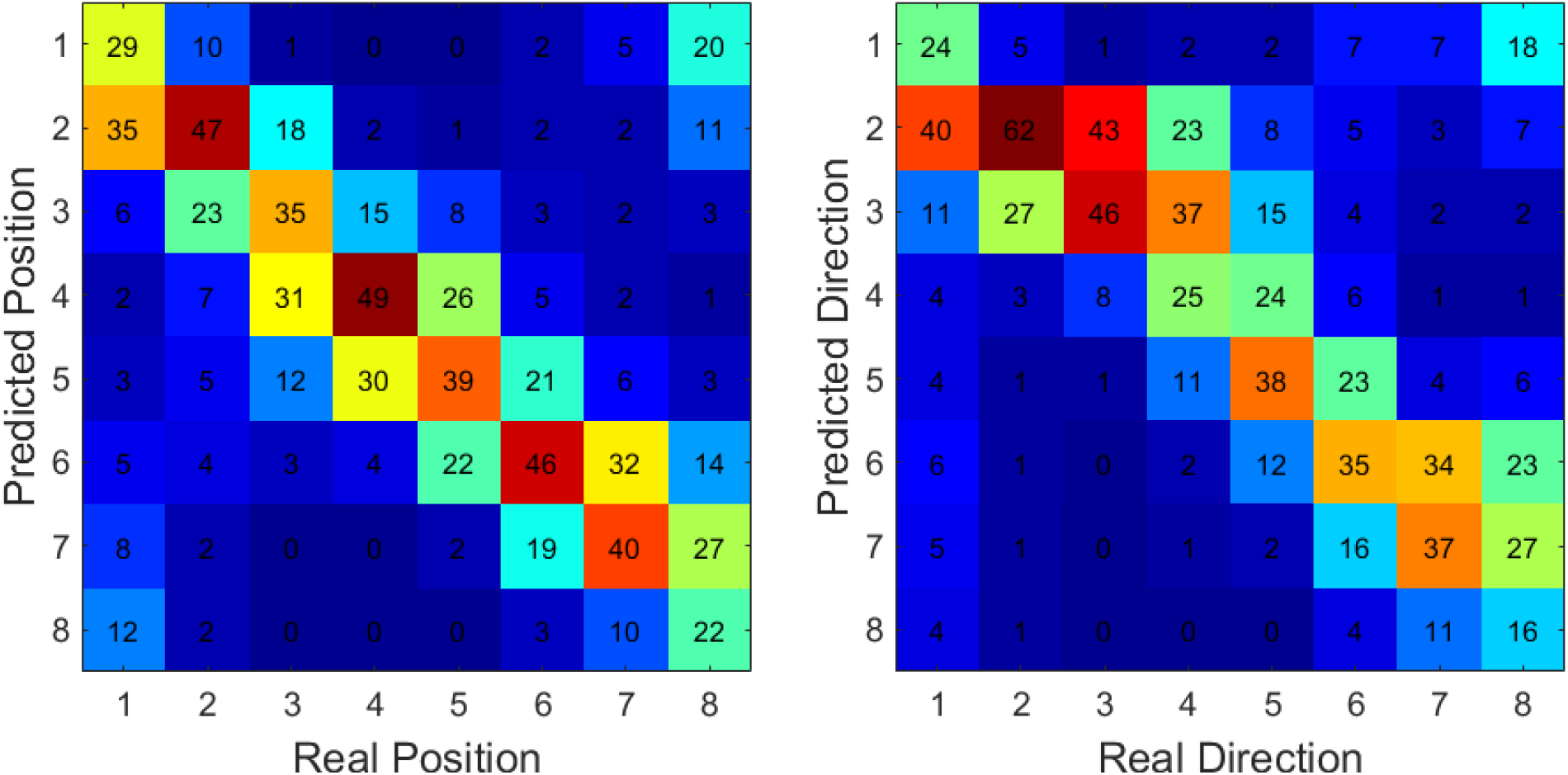
Confusion matrices for results shown in Figure 10a, b. The saccade end positions were predicted based on the activity of a sample of 25 neurons that have previously shown the strongest similarity between the activity profiles for regular and interceptive saccades. They were then assigned to one of the eight tested positions based on minimal Euclidian distance. For regular saccades (left) on average 38% of the trials were assigned correctly and in 82% correct or to one of the neighbors. For interceptive saccades (right) on average 35% of the trials were assigned correctly and in 78% correct or to one of the neighbors.

